# Genetic and epigenetic architectures of neurological protein biomarkers in the Lothian Birth Cohort 1936

**DOI:** 10.1101/558940

**Authors:** Robert F. Hillary, Daniel L. McCartney, Sarah E. Harris, Anna J. Stevenson, Anne Seeboth, Qian Zhang, David C. Liewald, Kathryn L. Evans, Craig W. Ritchie, Elliot M. Tucker-Drob, Naomi R. Wray, Allan F. McRae, Peter M. Visscher, Ian J. Deary, Riccardo E. Marioni

**Affiliations:** Centre for Genomic and Experimental Medicine, Institute of Genetics and Molecular Medicine, University of Edinburgh, Edinburgh, EH4 2XU; Centre for Cognitive Ageing and Cognitive Epidemiology, University of Edinburgh, Edinburgh, EH8 9JZ; Department of Psychology, University of Edinburgh, Edinburgh, EH8 9JZ; Institute for Molecular Bioscience, University of Queensland, Brisbane, QLD, Australia; Edinburgh Dementia Prevention, Centre for Clinical Brain Sciences, University of Edinburgh, Edinburgh, EH16 4UX; Department of Psychology, The University of Texas at Austin, United States; Population Research Center, The University of Texas at Austin, United States

## Abstract

Although plasma proteins may serve as important markers of disease risk in neurological conditions, the molecular mechanisms responsible for inter-individual variation in plasma protein levels are poorly understood. In this study, we conducted genome- and epigenome-wide association studies on the levels of 92 neurological proteins to identify genetic and epigenetic loci associated with their plasma concentrations (n = 750). We identified 62 independent genome-wide significant loci for 37 proteins (P < 5.4 × 10^−10^) and 68 epigenome-wide significant sites associated with the levels of 7 proteins (P < 3.9 × 10^−10^). Using this information, we identified biological pathways in which putative neurological biomarkers are implicated as well as molecular mechanisms through which genetic variation may perturb plasma protein levels. Additionally, we found evidence that poliovirus receptor is causally associated with Alzheimer’s disease. In conclusion, we identified many novel genetic and epigenetic factors that are associated with neurological protein levels which may inform disease biology and establish causal relationships between biomarkers and neurological diseases.

## 1. Introduction

Plasma proteins execute diverse biological processes and aberrant levels of these proteins are implicated in various disease states. Consequently, plasma proteins may serve as biomarkers, contributing to individual disease risk prediction and personalised clinical management strategies (Geyer *et al*., 2017). Identifying circulating biomarkers is of particular importance in neurological disease states in which access to diseased neural tissue *in vivo* is almost impossible. Furthermore, in neurodegenerative disorders, symptomatology may appear in only advanced clinical states, necessitating early detection and intervention (Polivka *et al*., 2016). Elucidating the factors which underpin inter-individual variation in plasma protein levels can inform disease biology and also identify proteins with likely causal roles in a given disease, augmenting their value as predictive biomarkers. Indeed, studies have characterised genetic variants (protein quantitative trait loci; pQTLs) associated with circulating protein levels and utilised such genetic information to identify proteins with causal roles in conditions such as cardiovascular diseases (Yao *et al*., 2018, Sun *et al*., 2018, Suhre *et al*., 2017). However, studies which have aimed to examine the genetic architecture of neurology-related proteins in human plasma are limited (Kim *et al*., 2011, Kauwe *et al*., 2014, Sasayama *et al*., 2017). Furthermore, few studies have combined genetic with epigenetic data to provide an additional layer of information regarding the molecular mechanisms responsible for regulating blood protein levels (Ahsan *et al*., 2017). Therefore, the goal of the present study was to characterise the genetic and epigenetic (using DNA methylation) architectures of putative neurology-related protein biomarkers in order to identify potential molecular determinants which regulate their plasma levels.

Here, genome- and epigenome-wide association studies (GWAS/EWAS) are carried out on the plasma levels of 92 neurological proteins in 750 participants from the Lothian Birth Cohort 1936 study (levels adjusted for age, sex, population structure and array plate; hereafter simply referred to as protein levels). These proteins represent the Olink® neurology panel and encompass a mixture of proteins with established links to neurobiological processes (such as axon guidance and synaptic function) and neurological diseases (such as Alzheimer’s disease (AD)), as well as exploratory proteins with roles in processes including cellular regulation, immunology and development. Following the identification of genotype-protein associations (pQTLs), functional enrichment analyses are performed on independent pQTL variants. Upon identification of epigenetic factors associated with protein levels, tissue specificity and pathway enrichment analyses are conducted to reveal possible biological pathways in which neurological proteins are implicated. Protein QTL data are integrated with publicly available expression QTL data to probe the molecular mechanisms which may modulate circulating protein levels. Finally, GWAS summary data for proteins and disease states are integrated using two-sample Mendelian Randomisation to determine whether selected proteins are causally associated with neurological disease states.

## 2. Results

### 2.1 Genetic architecture of neurological protein biomarkers

For the GWAS, a Bonferroni P value threshold of 5.4 × 10^−10^ (genome-wide significance level: 5.0 × 10^−8^/92 proteins) was set. The GWAS analysis in 750 older adults identified 2,734 significant SNPs associated with 37 proteins (*P* < 5.4 × 10^−10^) (Figure 1a; Supplementary Table 1). Pruning of pQTL variants (LD *r*^2^ < 0.1) using the SNP2GENE function in FUMA (FUnctional Mapping and Annotation analysis) yielded 62 independent variants (Supplementary Table 2). Of these 62 variants, 56 (90.3%) were *cis* pQTLs (SNP within 10 Mb of the transcription start site (TSS) of the gene) and 6 (9.7%) were *trans* variants. Furthermore, *cis* only associations were present for 31/37 proteins (83.8%), compared to *trans* only associations for 4/37 proteins (10.8%). Two proteins (5.4%) were associated with both *cis* and *trans* pQTLs (CD200R and Siglec-9). For all independent *cis* pQTLs associated with a given protein, the pQTL with the lowest P value was denoted as the sentinel variant (n = 33). The significance of *cis* associations decreased as the distance of the sentinel variant from the TSS increased (Figure 1b).

**Figure 1.**
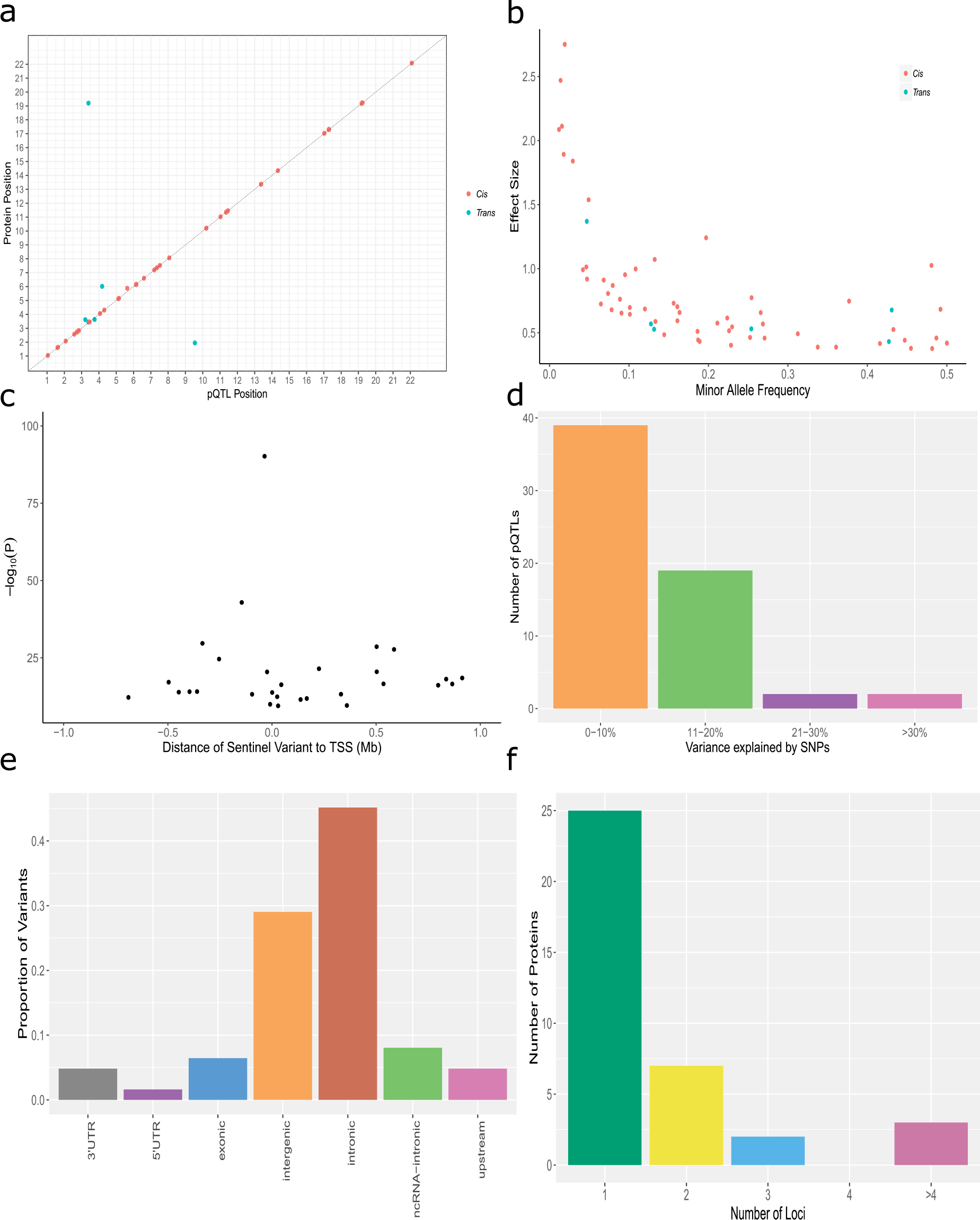
**(a)** Chromosomal locations of pQTLs. The x-axis represents the chromosomal location of independent *cis* and *trans* SNPs associated with the levels of Olink® neurology proteins. The y-axis represents the position of the gene encoding the associated protein. *Cis* (red); *trans* (blue). **(b)** Significance of sentinel *cis* variants versus distance of variants from the gene transcription start site. **(c)** Absolute effect size (per standard deviation of difference in protein level per effect allele) of independent pQTLs versus minor allele frequency. *Cis* (red); *trans* (blue). **(d)** Variance in protein levels explained by independent pQTLs. It is important to note that these estimates may be inflated owing to winner’s curse or over-fitting in the discovery GWAS **(e)** Classification of pQTL variants by function as defined by functional enrichment analysis in FUMA. **(f)** Number of loci significantly associated with independent pQTL variants per Olink® neurology protein.

The minor allele frequency of independent pQTL variants was inversely associated with effect size (Figure 1c). Notably, this association may be, in part, due to ascertainment bias as rarer variants (with lower minor allele frequencies) must have large effect sizes to attain the same level of power as more common variants. Independent pQTLs explained between 5.3% (rs2706505; CLM-6; P = 3.4 × 10^−10^) and 54.6% (rs9349050; MDGA1; P = 6.6 × 10^−91^) of the phenotypic variance in plasma protein levels (Supplementary Table 2; Figure 1d). The majority of pQTL variants were located in intergenic and intronic regions (Supplementary Table 2; Figure 1e). The number of independent loci associated per protein is shown in Figure 1f. No independent pQTL was associated with more than one protein. However, 28 non-independent *trans* pQTLs were associated with levels of Siglec-9 and CD200R. These variants were annotated to the *ST3GAL6-AS1* gene. Figure 2 demonstrates the effect of genetic variation at the most significant *cis* pQTL (rs9349050; MDGA1) and *trans* pQTL (rs12496730; Siglec-9) on protein levels.

**Figure 2.**
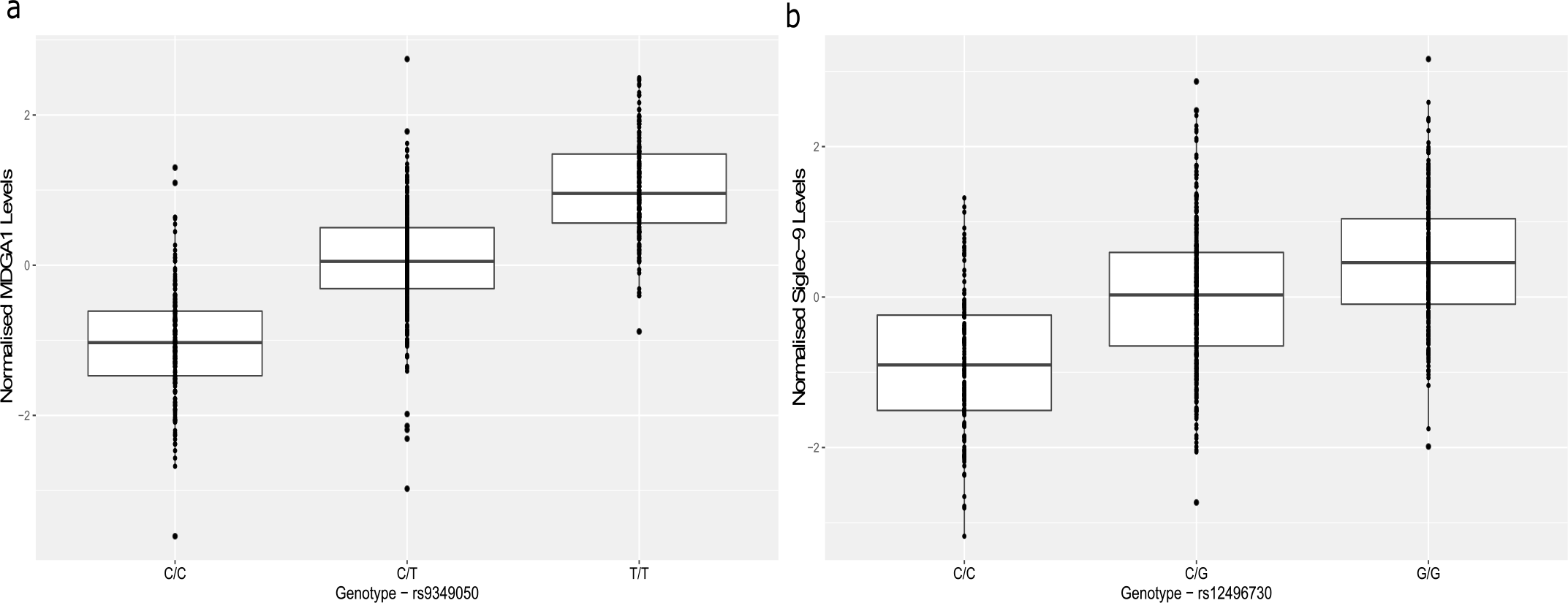
Effect of genetic variation on neurological protein levels. **(a)** Box plot of MDGA1 levels as a function of genotype (rs9349050, effect allele: T, other allele: C, beta = 1.03, se = 0.05). **(b)** Box plot of Siglec-9 levels as a function of genotype (rs12496730, effect allele: C, other allele: G, beta = −0.68, se = 0.06). Standard error (se).

We also used an alternative method, conditional and joint (COJO) analysis, to find independent pQTLs. This approach identified 40 significant pQTLs associated with the levels of 35 proteins (87.5% *cis* and 12.5% *trans* effects; Bonferroni-corrected level of significance: *P* < 5.4 × 10^−10^) (Supplementary Table 3). Seven independent pQTLs associated with the levels of 6 proteins were found using both approaches whereas the remaining SNPs identified by COJO for a given protein were located within the same locus as corresponding SNPs identified by FUMA.

### 2.2 Colocalisation of *cis* pQTLs with *cis* eQTLs

Of the 33 sentinel *cis* pQTL variants, 13 (39.3%) were *cis* eQTLs for the same gene in blood tissue. For 5/13 proteins, there was strong evidence (posterior probability (PP) > 0.75) for colocalisation of *cis* pQTLs and *cis* eQTLs and for 1 protein, NAAA, there was weaker evidence (PP > 0.5) for colocalisation. For 6/13 proteins, there was evidence (PP > 0.75) for two distinct causal variants affecting transcript and protein levels in the locus. Finally, for CLM-6, there was strong evidence (PP > 0.75) for a causal variant for gene expression only in the locus (Supplementary Table 4).

For the 5 proteins with strong evidence in favour of a shared causal variant for gene expression and plasma protein levels, two-sample Mendelian randomisation was performed to test for a causal association between perturbations in gene expression (using data from eQTLGen Consortium) and plasma protein levels (using our GWAS data). Pruned *cis* QTL variants (linkage disequilibrium (LD) r^2^ < 0.1) were used as instrumental variables for MR analyses. MR analyses were conducted using MR-Base (Hemani *et al*., 2018). For each trait, the intercept from MR Egger regression was non-significant, which does not suggest strong evidence for directional pleiotropy (DRAXIN: P = 0.82; Siglec-9: P = 0.41; MDGA1: P = 0.38; KYNU: P = 0.36; LAIR-2: P = 0.56). For 4 proteins, variation in gene expression was causally associated with plasma protein levels (Inverse variance-weighted method; Siglec-9: beta = 0.76, se = 0.26, P = 3.1 × 10^−3^; MDGA1: beta = 0.99, se = 0.49, P = 0.02; KYNU: beta = 1.05, se = 0.22, P = 2.2 × 10^−6^; LAIR-2: beta = 1.53, se = 0.51, P = 3.0 × 10^−3^). While gene expression of DRAXIN was not causally associated with changes in plasma protein levels (Inverse variance-weighted method; beta = −0.98, se = 0.62, P = 0.10), altered plasma protein levels were causally associated with changes in gene expression (beta = −0.72, se = 0.07, P = 1.2 × 10^−23^).

### 2.3 Epigenetic architecture of neurological protein biomarkers

For the EWAS, a Bonferroni P value threshold of 3.9 × 10^−10^ (genome-wide significance level: 3.6 × 10^−8^/92 proteins) was set (Saffari *et al*., 2018). We identified 68 genome-wide significant CpG sites associated with the levels of 7 neurological proteins (P < 3.9 × 10^−10^). Of these associations, 52 were *trans* effects (76.5%) and 16 associations were *cis* effects (23.5%) (Figure 3; Supplementary Table 5). The majority of the protein-CpG associations were attributable to CRTAM (Cytotoxic and Regulatory T Cell Molecule; 43/68; 63.2%; Figure 3), which is upregulated in CD4^+^ and CD8^+^ cells (Takeuchi *et al*., 2016).

**Figure 3.**
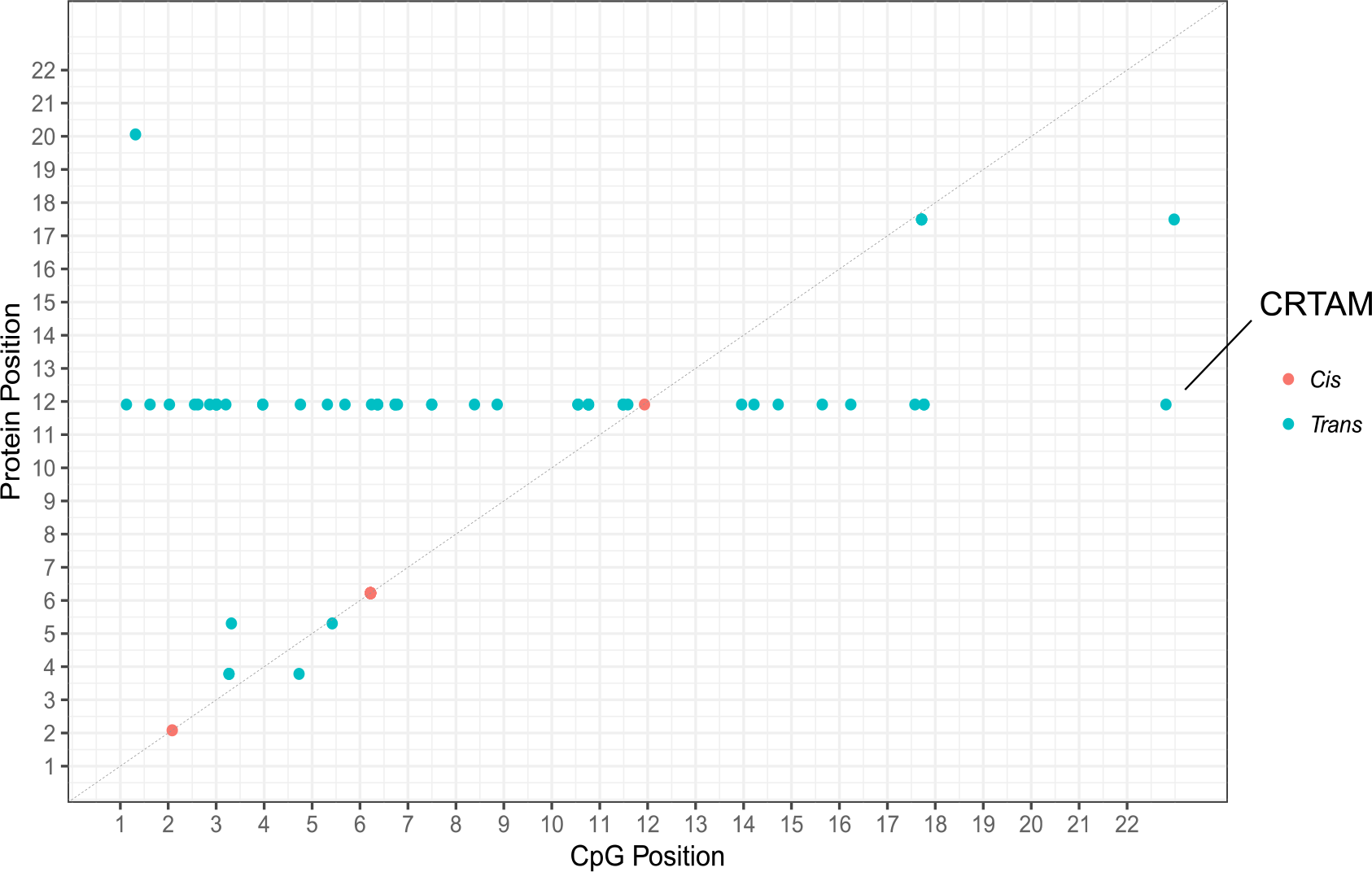
Genomic locations of CpG sites associated with differential neurological protein levels. The x-axis represents the chromosomal location of CpG sites associated with the levels of Olink® neurology biomarkers. The y-axis represents the position of the gene encoding the associated protein. Notably, *cis* CpG sites (n = 16) identified by our EWAS on protein levels lay within the same cluster for a given protein. These CpG sites lay too close to discriminate, resulting in the appearance of 3 CpG clusters in this figure. The protein on Chromosome 11 which constitutes the majority of methylome-protein associations is annotated (CRTAM). *Cis* (red); *trans* (blue).

Three proteins exhibited both genome-wide significant SNP and CpG site associations: MATN3, MDGA1 and NEP (Figure 4). For MATN3, the *cis* pQTL identified in this study (rs1147118) has previously been identified as a methylation QTL (mQTL) for the single *cis* CpG site associated with MATN3 levels identified by our EWAS (cg24416238) (Bonder *et al*., 2017). Similarly, the 6 *cis* pQTLs for differential blood MDGA1 concentrations in our study have been significantly associated with methylation levels of *cis* CpG sites identified by our EWAS on MDGA1 levels (Bonder *et al*., 2017). Finally, for NEP, we identified a sole independent *trans* pQTL (rs35004449) annotated to the *ITIH4* gene (beta: −0.53; effect allele: G) as well as three *trans* genome-wide significant CpG sites (cg18404041, cg11645453 and cg06690548 annotated to *ITIH1, ITIH4* and *SLC7A11,* respectively). In addition to lower circulating levels of NEP, this SNP has previously been associated with higher methylation levels of cg18404041 (*ITIH4*; beta: 0.93; effect allele: G; P = 4.20 × 10^−17^) and lower methylation levels of cg11645453 (*ITIH1*; beta: −0.84; effect allele: G; P = 1.20 × 10^−88^) (Gaunt *et al*., 2016).

**Figure 4.**
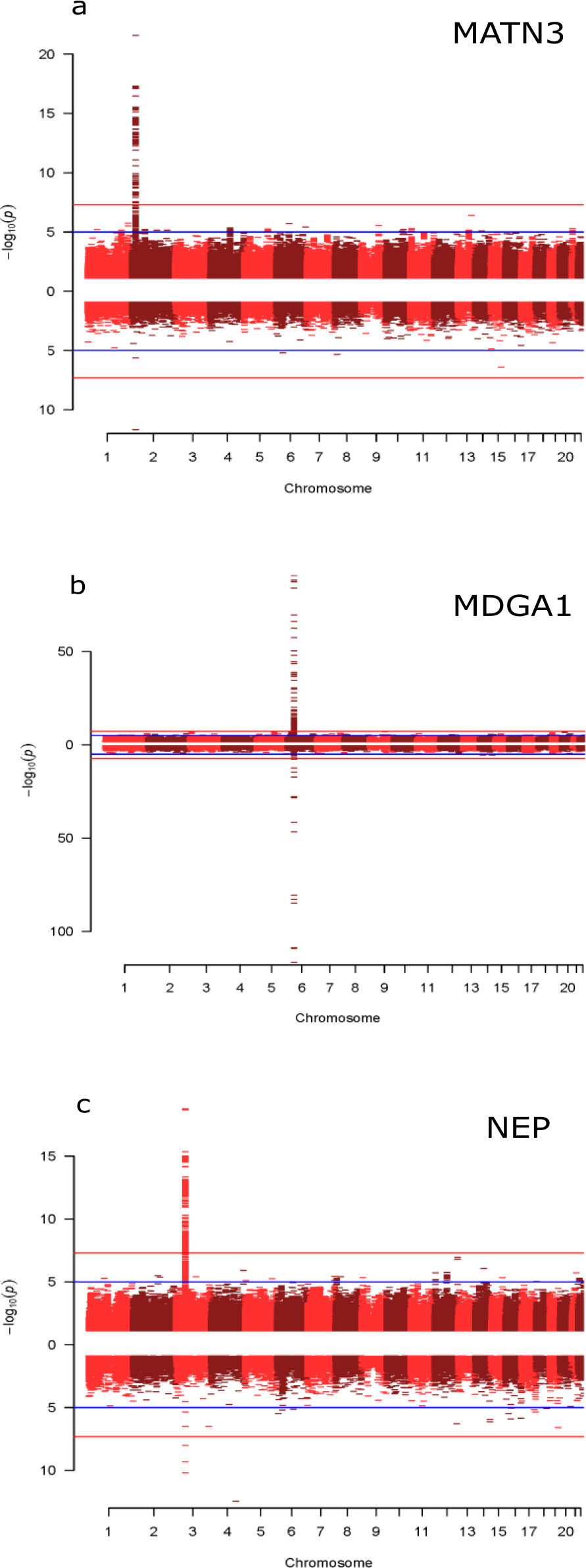
Miami plots of three neurological proteins with both genome-wide significant SNP and genome-wide significant CpG associations. The top half of the plot (skyline) shows the results from the GWAS on protein levels, whereas the bottom half (waterfront) shows the results from the EWAS. Blue lines indicate suggestive associations; red lines indicate epigenome-wide significant associations. **(a)** Miami plot for MATN3 (chromosome 2: 20,191,813-20,212,455). **(b)** Miami plot for MDGA1 (chromosome 6: 37,600,284-37,667,082). **(c)** Miami plot for NEP (chromosome 3: 154,741,913-154,901,518).

We conducted tissue specificity and pathway enrichment analyses (KEGG and GO – see methods for details) based on genes identified by methylation for each of the 7 proteins with genome-wide significant CpG associations. Tissue-specific patterns of expression were observed for 6/7 proteins (Supplementary Data 1). Neural tissue was the most common tissue type in which genes were differentially expressed (n = 5/6 proteins), followed by cardiac and splenic tissue (n = 4/6 proteins). Additionally, KEGG pathway analysis revealed that genes identified in the EWAS on CRTAM levels were enriched for neuroactive ligand-receptor interaction, axon guidance and cancer-related pathways (Supplementary Table 6; FDR-adjusted P value < 0.05). Gene ontology analyses revealed that genes incorporating CpG sites associated with CRTAM and GZMA levels are over-represented in neurogenesis, cell and neuronal differentiation, and synaptic organisation pathways (Supplementary Table 7-8; FDR-adjusted P value < 0.05). Furthermore, genes annotated to CpG sites associated with circulating SIGLEC1 and G-CSF levels are over-represented in immune system processes, viral response and cytokine response pathways (Supplementary Table 9-10; FDR-adjusted P value < 0.05). Finally, genes incorporating CpG sites associated with NEP levels are over-represented in metabolic pathways involving extracellular matrix components (Supplementary Table 11; FDR-adjusted P value < 0.05). For MDGA1 and MATN3, there were no significant results following multiple testing correction.

### 2.4 Correlation between genetic architecture of Olink neurology proteins and neurological phenotypes

Weak correlations were observed between all protein levels and 13 neurological phenotype polygenic risk scores (range: r = −0.11 to 0.11; Figure 5, Supplementary Table 12). Across all protein-polygenic risk score (PRS) comparisons, the most positive correlation was between CDH3 and the PRS for attention-deficit hyperactivity-disorder (r = 0.11, 95% CI: 0.04 – 0.19, P = 9.0 × 10^−4^). The most negative correlation was between NCAN and the PRS for depression as diagnosed by criteria stipulated by International Classification of Diseases 9/10 (r = −0.11, 95% CI: −0.04 – −0.18, P = 2.4 × 10^−3^). No relationship remained significant after correction for multiple testing (Bonferroni threshold: 0.05/(13 phenotypes x 92 proteins) = 0.05/1196 = 4.2 × 10^−5^).

**Figure 5.**
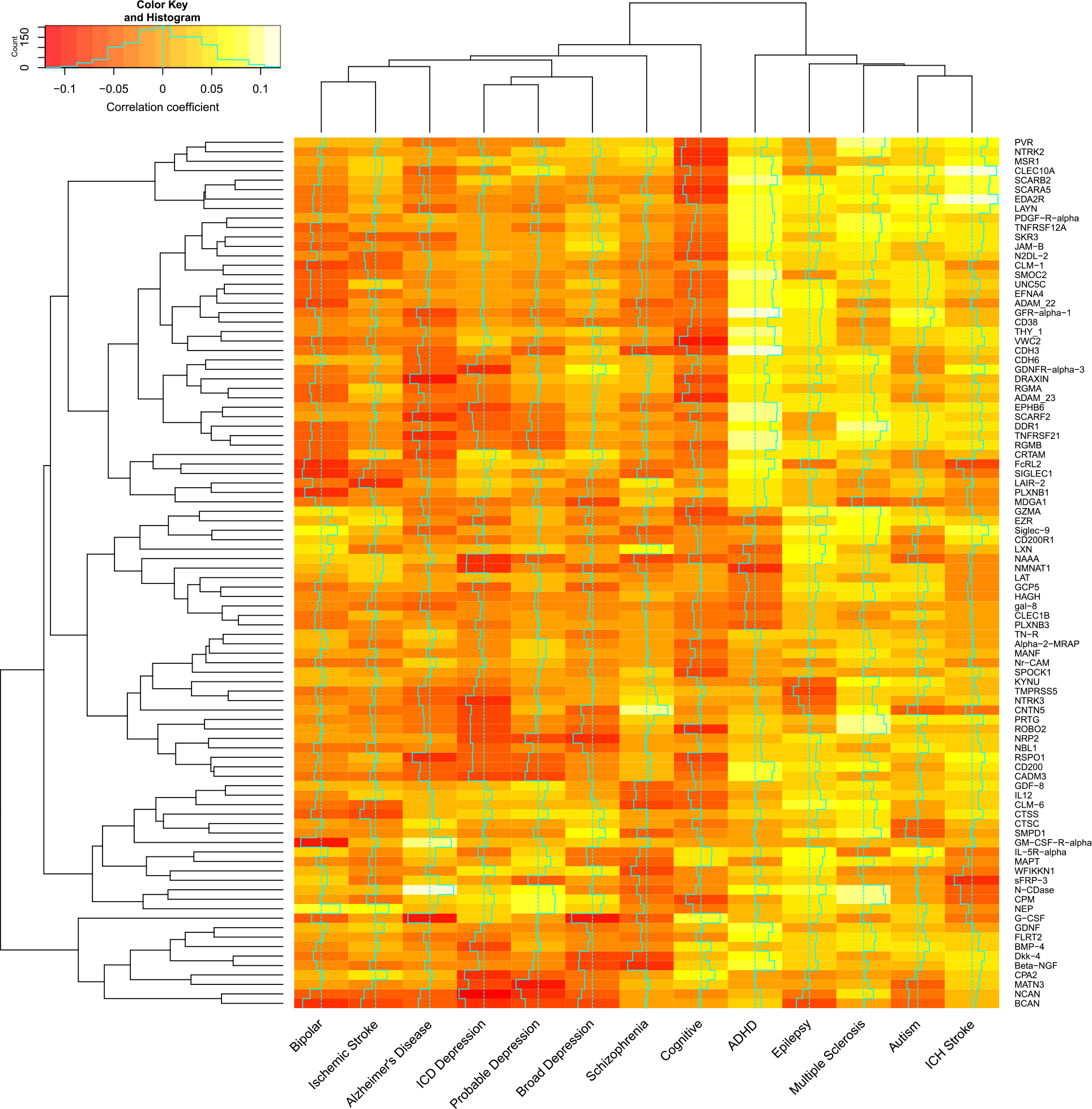
Correlation of polygenic risk scores for neurological phenotypes versus neurological protein levels. Weak correlations were observed between all polygenic risk scores and protein levels (r: −0.1 to 0.1). Red tiles indicate negative correlations whereas yellow tiles indicate positive correlations. ICD (International Classification of Diseases); ICH (Intracerebral Haemorrhage).

### 2.5 Causal evaluation of biomarkers in neurological disease

As the pQTL GWAS identified a *cis* association for poliovirus receptor (PVR), and given that the *PVR* gene has been implicated in AD (Porcellini *et al*., 2010), colocalisation analysis was performed to test if the same SNP variant might be driving both associations. A 200kb region surrounding the sentinel *cis* pQTL for PVR was extracted from GWAS summary statistics for PVR levels as well as AD (Jansen *et al*., 2019). Default priors were applied. There was evidence to suggest that there are two distinct causal variants for altered protein levels and AD risk within the region (PP3 > 0.99).

In addition to the colocalisation analysis, two-sample Mendelian randomisation was used to test for putatively causal associations between plasma PVR levels and AD (Lambert *et al*., 2013). After LD pruning, only one independent SNP remained (rs7255066; F-statistic: 15.88). Therefore, causal effect estimates were determined using the Wald ratio test, i.e., a ratio of effect per risk allele on AD to effect per risk allele on PVR levels. MR analyses indicated that PVR levels were causally associated with AD (beta = 0.17, se = 0.02, P = 5.2 × 10^−10^; Wald ratio test). Testing for horizontal pleiotropy was not possible owing to an insufficient number of instruments. Conversely, AD risk was not causally associated with PVR levels (number of SNPs: 5; Inverse variance-weighted method: beta = 0.38, se = 0.29, P = 0.34). The intercept from MR Egger regression was −0.08 (se: 0.08; P = 0.42) which does not provide strong evidence for directional pleiotropy.

### 2.6 Replication of previous pQTL studies

Replication of the pQTL findings was carried out via lookup of genotype-protein summary statistics from existing pQTL studies (Sun *et al*., 2018, Suhre *et al*., 2017, Di Narzo *et al*., 2017, Lourdusamy *et al*., 2012). Of the 37 proteins with a pQTL in the present study, 16 (with 24 QTLs) were available for lookup. In total, 9/23 (39.1%) pQTLs replicated at P < 1.25 × 10^−7^ (denoting the least conservative threshold across all studies) (Supplementary Table 13).

## 3. Discussion

Using a multi-omics approach, we identified 62 independent genome-wide significant pQTLs and 68 genome-wide significant CpG sites associated with circulating neurological protein levels. To probe the molecular mechanisms which modulate plasma protein levels, we integrated pQTL and eQTL data allowing for the examination of whether pQTLs affect gene expression. For five proteins, we found strong evidence that a common causal variant underpinned changes in transcript and protein levels. Mendelian randomisation analyses suggested that variants for four of these proteins (Siglec-9, MDGA1, KYNU and LAIR-2) influence protein levels by altering gene expression. However, for one protein (DRAXIN), the converse may be true as our data suggested that altered plasma protein levels of this neurodevelopmental protein may affect gene expression, perhaps through a feedback mechanism. Genotype-protein associations for other proteins may exert their influence on protein levels through modulation of protein clearance, degradation, binding or secretion. Methylation data revealed that neurological proteins were implicated in immune, neurological development and synaptic functionality pathways. Finally, we found evidence of a causal association between poliovirus receptor protein and AD.

In addition to leveraging methylation data to identify pathway enrichment for plasma proteins, identification of *trans* pQTLs may highlight previously unidentified pathways relevant to disease processes. For instance, we found that genetic variation at the inter-alpha-trypsin inhibitor heavy chain family member 4 locus (*ITIH4)* is associated with differential NEP levels (*trans* pQTL: rs35004449). Additionally, two CpG sites annotated to *ITIH4* and *ITHI1* (cg18404041 and cg11645453, respectively) were associated with NEP levels. Methylation QTL analyses revealed that the SNP rs35004449 has been previously associated with higher methylation levels of cg18404041 (*ITIH4*) and lower DNA methylation levels of cg11645453 (*ITIH1*) (Gaunt *et al*., 2016). Similarly, this SNP has been associated with lower gene expression of *ITIH4* (Consortium, 2015) and higher protein levels of *ITIH1* (Sun *et al*., 2018). Together, these data suggests that the expression of NEP, ITIH4 and ITIH1 may be co-regulated, involving inverse relationships between NEP and ITIH4 with ITIH1. Given that mutations in *NEP* have been linked to Alzheimer’s pathology and that upregulation of ITIH4 has been demonstrated in sera of AD patients (Yang *et al*., 2012b), mechanistic studies relating to co-expression of these proteins are merited in pathological states.

In this study, no independent pQTL was associated with more than one protein. However, prior to pruning of SNPs to identify independent signals, 28 SNPs were shared between CD200R and Siglec-9. These polymorphisms mapped to the *ST3GAL6-AS1* gene and included the sentinel SNP for Siglec-9, but not CD200R1. ST3GAL6-AS1 is a long-coding RNA which is associated with increased expression of ST3GAL6, an enzyme responsible for catalysing the addition of sialic acid to cell surfaces (Bull *et al*., 2014). Upregulation of ST3GAL6 has been reported in multiple myeloma (Shen *et al*., 2018, Glavey *et al*., 2014); this permits evasion of immune responses against cancer cells through binding of sialic acid to Siglec receptor proteins, such as Siglec-9. The recognition of sialic acid by Siglec proteins ignites signalling cascades which promotes immune inhibitory responses (Jandus *et al*., 2014, Adams *et al*., 2018). Furthermore, CD200-CD200R interaction results in the inhibition of immune responses against multiple myeloma cells (Conticello *et al*., 2013). Therefore, as polymorphisms in *ST3GAL6-AS1* are associated with altered expression of Siglec-9 and CD200R, this may provide further evidence for co-regulation of these proteins in pathological milieux, such as tumorigenesis in cancers including multiple myeloma. Polymorphisms in such *trans* pQTLs may also be used to predict disease risk, progression and provide pharmacogenomic information in predicting individual patient responses to inhibition of these co-regulated proteins.

By using *cis* pQTLs as instruments for MR analyses, it is possible test whether plasma proteins are associated causally with disease states (Zheng *et al*., 2017). *PVR* is a component of the AD risk-associated *APOE*/*TOMM40* cluster on chromosome 19 and has been hypothesised to influence risk of AD through susceptibility to viral infections (Porcellini *et al*., 2010). However, it is unknown whether PVR is causally linked to the disease. MR analyses suggested that circulating PVR levels may be causally associated with AD and not vice versa. However, an insufficient number of instruments were available to permit testing for potential pleiotropic effects. Furthermore, colocalisation analysis revealed that independent variants in the *PVR* locus are likely causally associated with altered plasma PVR levels and AD risk. One possible explanation for this is that variation at a pQTL is associated with circulating PVR levels but may not reflect disease-relevant mechanisms, such as altered *PVR* expression in neural tissue. Indeed, genetic variation at a distinct site in the locus may directly influence AD risk through a distinct neuropathological mechanism leading to development of the disease. Further studies are merited to define possible pleiotropic effects, as well as the precise relationship between genetic variation in the *PVR* locus and susceptibility to AD to refine its potential as a biomarker for the disease.

The discrepancy in replication of pQTLs reported in previous studies may be due to a number of factors. Firstly, the sample sizes of these studies (n < 100 (Lourdusamy *et al*., 2012, Di Narzo *et al*., 2017); n > 1,000 (Suhre *et al*., 2017, Sun *et al*., 2018)) are different from that of the present study (n = 750) leading to differences in statistical power. Secondly, diverse proteomic platforms may result in the detection of different genotype-protein associations. Depending on platform technology, susceptibility to cross-reactive events and detection of proteins in their free, versus complexed, forms can result in inappropriate readouts. SOMAmer technology, employed in the previous pQTL studies is a highly sensitive, aptamer-based platform which overcomes limitations associated with antibody-based methods, such as cross-reactivity (Gold *et al*., 2010). Moreover, Olink® technology is particularly effective in limiting the reporting of cross-reactive events. However, when compared to these platforms, other technologies such as mass-spectrometry can produce highly accurate measurements but with low sensitivity (Hathout, 2015). Lack of standardisation amongst proteomic platforms, insufficient power to detect associations and differences in study demographics may all contribute to variability in the detection of pQTLs for a given protein. Additionally, we performed both FUMA (LD-based method) and COJO (stepwise conditional regression) to identify independent pQTL-protein associations and found a small overlap (17%) between SNPs identified by both methods. However, SNPs which were differentially identified by COJO and FUMA for a given protein were located within the same region. Indeed, the maximum distance between discordant SNPs for a given protein was 3 Mb.

We acknowledge several limitations in the present study. Firstly, analyses were restricted to individuals of European descent, complicating the generalisability of our findings to individuals of other ethnic backgrounds. Secondly, functional enrichment analyses indicated that a number of *cis* pQTL variants alter the amino acid sequence of the coded protein which may impact the quantitative protein assay. Finally, as our findings pertain to whole blood samples, studies examining the genetic and epigenetic regulation of neurological proteins in *post-mortem* brain tissue are warranted.

In conclusion, we have identified genetic and epigenetic factors associated with neurological proteins in an older-age population. We have shown that use of a multi-omics approach can help define whether such proteins are causal in disease processes. Importantly, we have shown that PVR may be causally associated with AD. Furthermore, we have provided a platform upon which future studies can interrogate pathophysiological mechanisms underlying neurological conditions. Together, this information may help inform disease biology as well as aid in the prediction of disease risk and progression in clinical settings.

## 4. Online Methods

### 4.1 The Lothian Birth Cohort 1936

The Lothian Birth Cohort 1936 (LBC1936) comprises Scottish individuals born in 1936, most of whom took part in the Scottish Mental Survey 1947 at age 11. Participants who were living within Edinburgh and the Lothians were re-contacted approximately 60 years later, 1,091 consented and joined the LBC1936. Upon recruitment, participants were approximately 70 years of age (mean age: 69.6 ± 0.8 years). Participants subsequently attended four additional waves of clinical examinations every three years. Detailed genetic, epigenetic, physical, psychosocial, cognitive, health and lifestyle data are available for members of the LBC1936. Recruitment and testing of the LBC1936 cohort have been described previously (Deary *et al*., 2007, Taylor *et al*., 2018).

### 4.2 Protein Measurements in the Lothian Birth Cohort 1936

Plasma was extracted from 816 blood samples collected in citrate tubes at mean age 72.5 ± 0.7 years (Wave 2). Plasma samples were analysed using a 92-plex proximity extension assay (Olink® Bioscience, Uppsala Sweden). The proteins assayed constitute the Olink® neurology biomarker panel. In brief, 1 µL of sample was incubated in the presence of proximity antibody pairs linked to DNA reporter molecules. Upon binding of an antibody pair to their corresponding antigen, the respective DNA tails form an amplicon by proximity extension, which can be quantified by high-throughput real-time PCR. This method limits the reporting of cross-reactive events. The data were pre-processed by Olink® using NPX Manager software. Protein levels were transformed by rank-based inverse normalisation. Normalised plasma protein levels were then regressed onto age, sex, four genetic principal components of ancestry derived from the Illumina 610-Quadv1 genotype array (to control for population structure) and Olink® array plate. Standardised residuals from these linear regression models were used in our genome- and epigenome-wide association studies.

### 4.3 Methylation preparation in the Lothian Birth Cohort 1936

DNA from whole blood was assessed using the Illumina 450K methylation array at the Edinburgh Clinical Research Facility (Wave 2; n = 895; mean age: 72.5 ± 0.7 years). Details of quality control procedures have been described elsewhere (Shah *et al*., 2014). Briefly, raw intensity data were background-corrected and normalised using internal controls. Following background correction, manual inspection permitted removal of low quality samples presenting issues relating to bisulphite conversion, staining signal, inadequate hybridisation or nucleotide extension. Quality control analyses were performed to remove probes with low detection rate i.e. <95% at P < 0.01. Samples with a low call rate (samples with < 450,000 probes detected at p-values of less than 0.01) were also eliminated. Furthermore, samples were removed if they had a poor match between genotype and SNP control probes, or incorrect DNA methylation-predicted sex.

### 4.4 Genotyping in the Lothian Birth Cohort 1936

LBC1936 DNA samples were genotyped at the Edinburgh Clinical Research Facility using the Illumina 610-Quadv1 array (Wave 2; n = 1,005; mean age: 72.5 ± 0.7 years; San Diego). Preparation and quality control steps have been reported previously (Davies *et al*., 2011). Individuals were excluded on the basis of sex discrepancies, relatedness, SNP call rate of less than 0.95, and evidence of non-Caucasian descent. SNPs with a call rate of greater than 0.98, minor allele frequency in excess of 0.01, and Hardy-Weinberg equilibrium test with P ≥ 0.001 were included in analyses.

### 4.5 Polygenic risk scoring in the Lothian Birth Cohort 1936

Polygenic risk scores for 13 psychiatric and neurological traits were calculated using the PRSice software program with LD clumping parameters set to R^2^ > 0.25 over 250 kilobase sliding windows (Supplementary Table 14) (Euesden *et al*., 2015). Polygenic scores were constructed using all SNPs (P < 1) from discovery GWAS (Davies *et al*., 2018, Sawcer *et al*., 2011, Schizophrenia Working Group of the Psychiatric Genomics, 2014, Howard *et al*., 2018, Cross-Disorder Group of the Psychiatric Genomics, 2013, Woo *et al*., 2014, International League Against Epilepsy Consortium on Complex Epilepsies. Electronic address, 2014, Jansen *et al*., 2019).

### 4.6 Genome-wide association studies

SNPs were imputed to the 1000G reference panel (phase 1, version 3). Genome-wide association analyses were conducted on 8,683,751 autosomal variants against protein residuals in 750 individuals from the Lothian Birth Cohort 1936. Linear regression was used to assess the effect of each genetic variant on the protein residuals using mach2qtl (Li *et al*., 2009, Li *et al*., 2010).

### 4.7 Epigenome-wide association studies

Epigenome-wide association analyses were conducted by regressing each of 459,309 CpG sites on transformed protein levels using linear regression with adjustments for age, sex, measured white blood cell counts (basophils, eosinophils, neutrophils, lymphocytes, monocytes) and technical covariates (plate, position, array, hybridisation, date). Outliers for white blood cell counts (n = 22) were excluded prior to analyses. Complete methylation and proteomic data were available for 692 individuals. Genome-wide significant CpG associations mapped to sites with underlying polymorphisms were excluded, as well as those predicted to cross-hybridise based on findings by Chen *et al*. (Chen *et al*., 2013). Analyses were performed using the limma package in R (Ritchie *et al*., 2015).

Pathway enrichment was assessed among KEGG pathways and Gene Ontology (GO) terms via hypergeometric tests using the *phyper* function in R. Furthermore, tissue specificity analyses were conducted using the GENE2FUNC function in FUnctional Mapping and Annotation (FUMA; outlined in the next section). Differentially expressed gene sets with Bonferroni-corrected P values of < 0.05 and an absolute log-fold change of ≥ 0.58 (default settings) were considered to be enriched in a given tissue type (GTEx v7).

### 4.8 Functional mapping and annotation of pQTLs

The identification of independent pQTL variants from the GWAS which yielded significant genotype-protein associations, and their subsequent functional annotation, were performed using the independent SNP algorithm implemented FUMA analysis (Watanabe *et al*., 2017). Initial independent significant SNPs were identified using the SNP2GENE function. These were defined as variants with a P value of < 5 × 10^−8^ that were independent of other genome-wide significant SNPs at r^2^ < 0.6. Lead independent SNPs, brought forward for this study, were further defined as the initial independent significant SNPs that were independent from each other at r^2^ < 0.1. Independent significant SNPs were functionally annotated using ANNOVAR (Wang *et al*., 2010) and Ensembl genes (build 85).

### 4.9 Conditional analysis

In addition to FUMA, we performed approximate genome-wide stepwise conditional analysis through GCTA-COJO using the ‘cojo-slct’ option in order to identify independent associations (Yang *et al*., 2012a). Conditional analyses were run per chromosome or per locus with the default settings of the software.

### 4.10 Characterisation of *cis* and *trans* effects

Genome-wide significant pQTLs and CpG sites were categorised into *cis* and *trans* effects. *Cis* associations were defined as loci which reside within 10 Mb of the TSS of the gene encoding the protein of interest. *Trans* effects were defined as those loci which lay outside of this region or were located on a chromosome distinct from that which harboured the gene TSS. TSS positions were defined using the biomaRt package in R (Durinck *et al*., 2009, Durinck *et al*., 2005) and Ensembl v83.

### 4.11 Identification of overlap between *cis* pQTLs and eQTLs

We cross-referenced sentinel *cis* pQTLs with publicly available *cis* eQTL data from the eQTLGen consortium (Võsa *et al*., 2018). *Cis* eQTLs were filtered to retain only variants with P < 5.4 × 10^−10^. Furthermore, only *cis* eQTLs for the same gene as the *cis* pQTL protein were retained. These associations were then tested for colocalisation.

### 4.12 Colocalisation analysis

To test the hypothesis that a single causal variant might underlie both an eQTL and pQTL, resulting in modulation of transcript and protein levels, we conducted Bayesian tests of colocalisation. Colocalisation analyses were performed using the coloc package in R (Giambartolomei *et al*., 2014). For each pQTL variant, a 200 kb region (upstream and downstream) was extracted from our GWAS summary statistics for each protein of interest. This window previously has been recommended in order to capture *cis* eQTLs, which often lie within 100 kb of their target gene (Guo *et al*., 2015). Expression QTLs for genes within this region were extracted from eQTLGen consortium summary statistics and subset to the gene encoding the protein of interest (Võsa *et al*., 2018). All SNPs shared by transcripts and proteins were used to determine the posterior probability for five distinct hypothesis. Default priors were applied. Posterior probabilities (PP) > 0.75 provided strong evidence in favour of a given hypothesis. Hypothesis 4 states that two association signals were attributable to the same causal variant. Associations with PP4 > 0.75 were deemed highly likely to colocalise. Associations with PP3 > 0.75 provided strong evidence for hypothesis 3 that there were independent causal variants for protein level and gene expression. In this study, hypothesis 2 referred to a causal variant for condition 2 (gene expression only) whereas hypothesis 1 represented a causal variant for protein levels only. Associations with PP0 > 0.75 (for hypothesis 0) indicated that it is highly likely there were no causal variants for either trait in the region.

### 4.13 Ethical approval

Ethical permission for the LBC1936 was obtained from the Multi-Centre Research Ethics Committee for Scotland (MREC/01/0/56) and the Lothian Research Ethics Committee (LREC/2003/2/29). Written informed consent was obtained from all participants.

## Acknowledgements

The authors thank all LBC1936 study participants and research team members who have contributed, and continue to contribute, to ongoing LBC1936 studies. The LBC1936 is supported by Age UK (Disconnected Mind program) and the Medical Research Council (MR/M01311/1). Methylation typing was supported by Centre for Cognitive Ageing and Cognitive Epidemiology (Pilot Fund award), Age UK, The Wellcome Trust Institutional Strategic Support Fund, The University of Edinburgh, and The University of Queensland. This work was conducted in the Centre for Cognitive Ageing and Cognitive Epidemiology, which is supported by the Medical Research Council and Biotechnology and Biological Sciences Research Council (MR/K026992/1), and which supports IJD. We acknowledge NIH Grants R01AG054628 and R01AG05462802S1 for supporting this research and Grant P2CHD042849 for supporting the Population Research Center at the University of Texas. RFH and AJS are supported by funding from the Wellcome Trust 4-year PhD in Translational Neuroscience – training the next generation of basic neuroscientists to embrace clinical research [108890/Z/15/Z]. AS is supported by a Medical Research Council PhD Studentship in Precision Medicine with funding by the Medical Research Council Doctoral Training Programme and the University of Edinburgh College of Medicine and Veterinary Medicine. DLMcC and REM are supported by Alzheimer’s Research UK major project grant ARUK-PG2017B-10. This research was supported by Australian National Health and Medical Research Council (grants 1010374, 1046880, and 1113400) and by the Australian Research Council (DP160102400). PMV, NRW, and AFM are supported by the NHMRC Fellowship Scheme (1078037, 1078901, and 1083656).

## Author’s Contributions

Conception and design: RFH and REM. Data analysis: RFH, DLMcC, DCL and REM. Drafting the article: RFH and REM. Revision of the article: all authors.

