## Supplementary Data 1 for "Genetic and epigenetic architectures of neurological protein biomarkers in the Lothian Birth Cohort 1936"

**Supplementary Data 1.** Tissue-specific patterns of expression for genes associated with Olink® neurology proteins exhibiting genome-wide significant CpG sites. Differentially expressed gene sets in 53 tissue types (GTEx v7) with Bonferroni-corrected P values of < 0.05 and an absolute log fold change ≥ 0.58 are shown in red. Differentially expressed genes (DEG).

### *CRTAM*

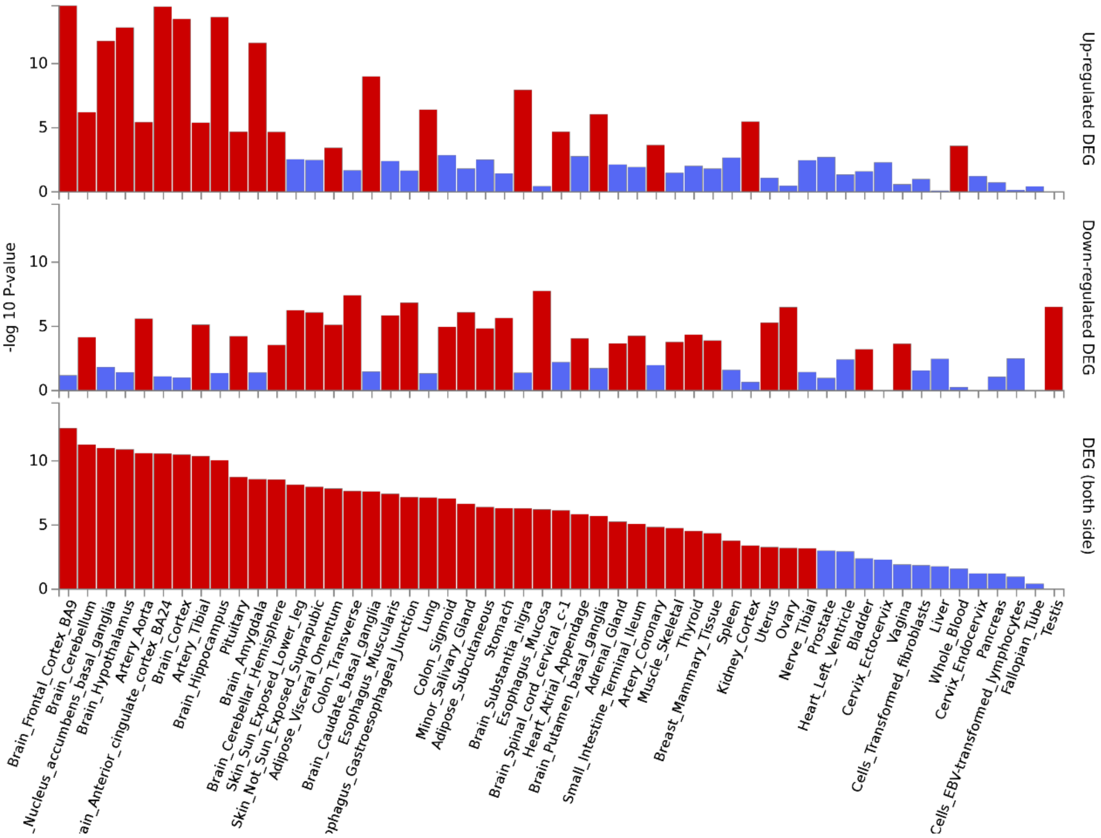

*G-CSF*

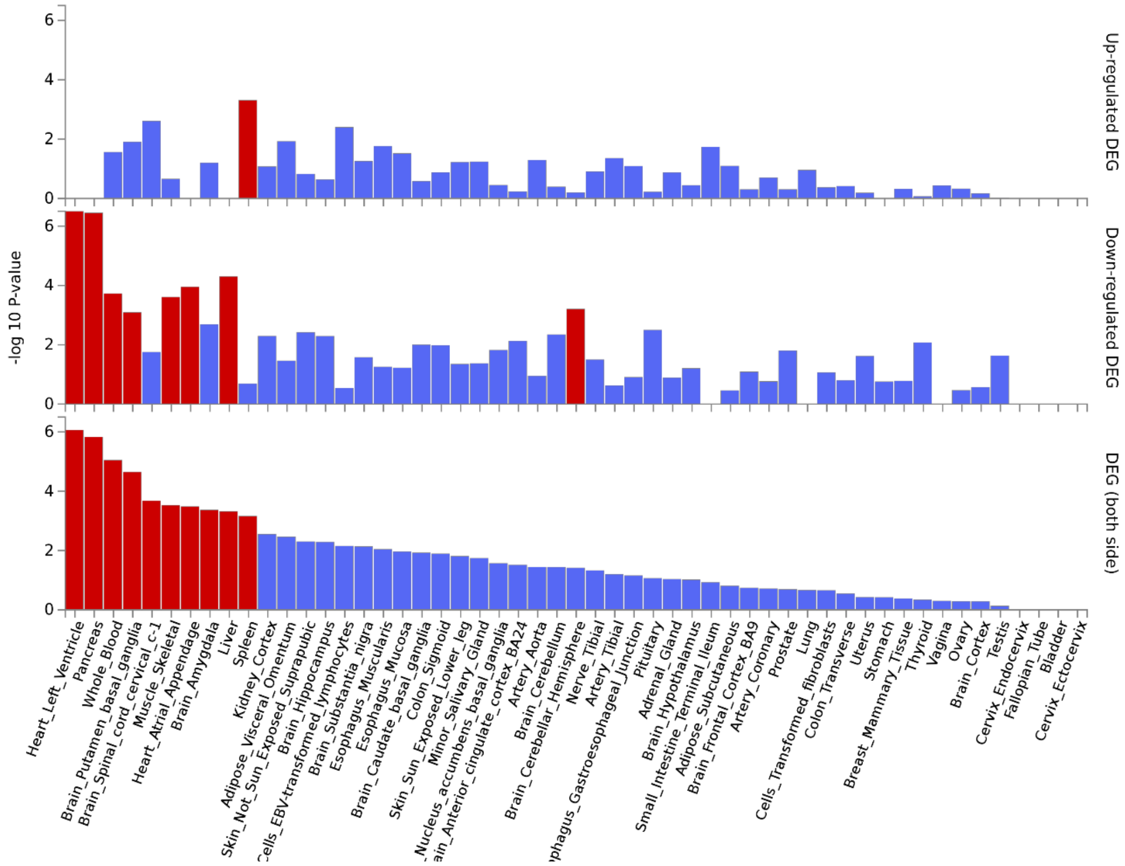

Up-regulated DEG
Down-regulated DEG
DEG (both side)
-log 10 P-value
0
2
4
6
Heart_Left_Ventricle
Pancreas
Whole_Blood
Brain_Putamen_basal_ganglia
Brain_Spinal_cord_cervical_c-1
Muscle_Skeletal
Heart_Atrial_Appendage
Brain_Amygdala
Liver
Spleen
Kidney_Cortex
Adipose_Visceral_Omentum
Skin_Not_Sun_Exposed_Suprapubic
Brain_Hippocampus
ells_EBV-transformed_lymphocytes
Brain_Substantia_nigra
Esophagus_Muscularis
Esophagus_Mucosa
Brain_Caudate_basal_ganglia
Colon_Sigmoid
Skin_Sun_Exposed_Lower_leg
Minor_Salivary_Gland
_Nucleus_accumbens_basal_ganglia
ain_Anterior_cingulate_cortex_BA24
Artery_Aorta
Brain_Cerebellum
Brain_Cerebellar_Hemisphere
Nerve_Tibial
Artery_Tibial
phagus_Gastroesophageal_Junction
Pituitary
Adrenal_Gland
Brain_Hypothalamus
Small_Intestine_Terminal_Ileum
Adipose_Subcutaneous
Brain_Frontal_Cortex_BA9
Artery_Coronary
Prostate
Lung
Cells_Transformed_fibroblasts
Colon_Transverse
Uterus
Stomach
Breast_Mammary_Tissue
Thyroid
Vagina
Ovary
Brain_Cortex
Testis
Cervix_Endocervix
Fallopian_Tube
Bladder
Cervix_Ectocervix

*GZMA*

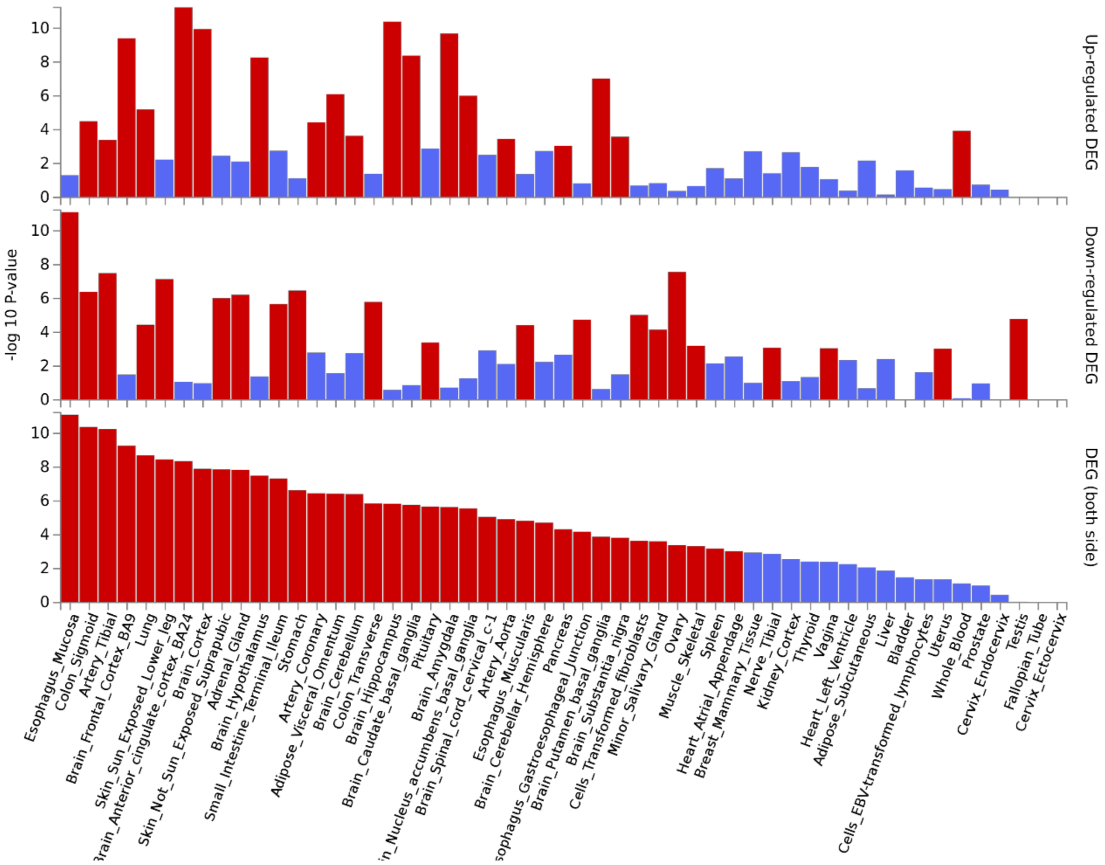

-log 10 P-value
Up-regulated DEG
Down-regulated DEG
DEG (both side)
Esophagus_Mucosa
Colon_Sigmoid
Artery_Tibial
Brain_Frontal_Cortex_BA9
Lung
Skin_Sun_Exposed_Lower_leg
Brain_Cortex
Skin_Not_Sun_Exposed_Suprapubic
Adrenal_Gland
Brain_Hypothalamus
Small_Intestine_Terminal_Ileum
Stomach
Artery_Coronary
Adipose_Visceral_Omentum
Brain_Cerebellum
Colon_Transverse
Brain_Hippocampus
Brain_Caudate_basal_ganglia
Pituitary
Brain_Amygdala
Brain_Spinal_cord_cervical_c-1
Artery_Aorta
Esophagus_Muscularis
Brain_Cerebellar_Hemisphere
Pancreas
Brain_Putamen_basal_ganglia
Brain_Substantia_nigra
Cells_Transformed_fibroblasts
Minor_Salivary_Gland
Ovary
Muscle_Skeletal
Spleen
Heart_Atrial_Appendage
Breast_Mammary_Tissue
Nerve_Tibial
Kidney_Cortex
Thyroid
Vagina
Heart_Left_Ventricle
Adipose_Subcutaneous
Liver
Bladder
Cells_EBV-transformed_lymphocytes
Uterus
Whole_Blood
Prostate
Cervix_Endocervix
Testis
Fallopian_Tube
Cervix_Ectocervix

*MDGA1*

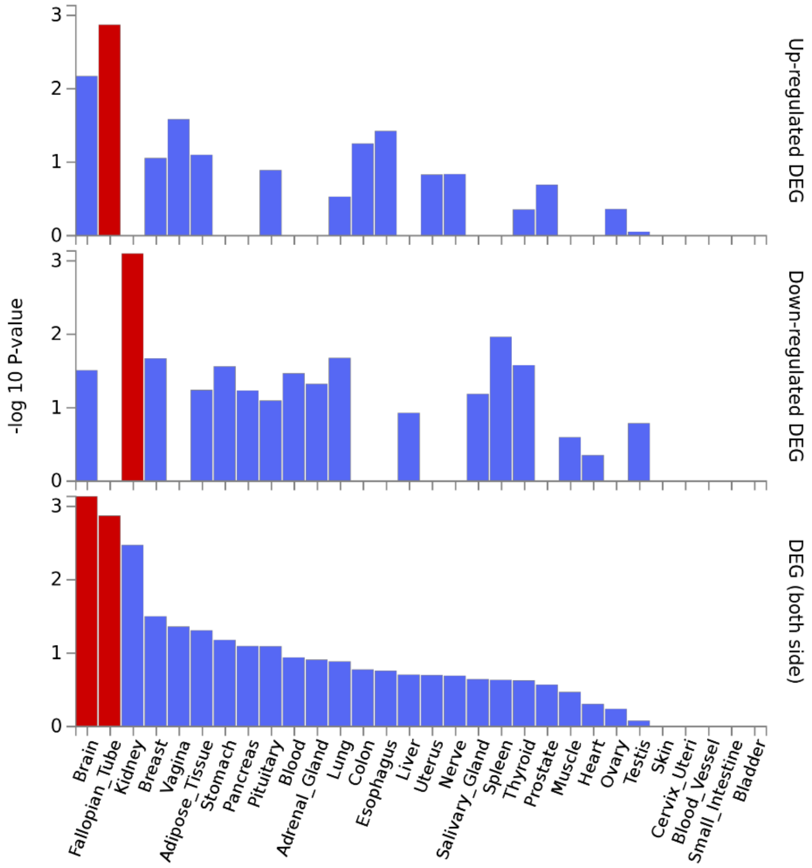

-log 10 P-value
Up-regulated DEG
Down-regulated DEG
DEG (both side)
0
1
2
3
Brain
Fallopian_Tube
Kidney
Breast
Vagina
Adipose_Tissue
Stomach
Pancreas
Pituitary
Blood
Adrenal_Gland
Lung
Colon
Esophagus
Liver
Uterus
Nerve
Salivary_Gland
Spleen
Thyroid
Prostate
Muscle
Heart
Ovary
Testis
Skin
Cervix_Uteri
Blood_Vessel
Small_Intestine
Bladder

*NEP*

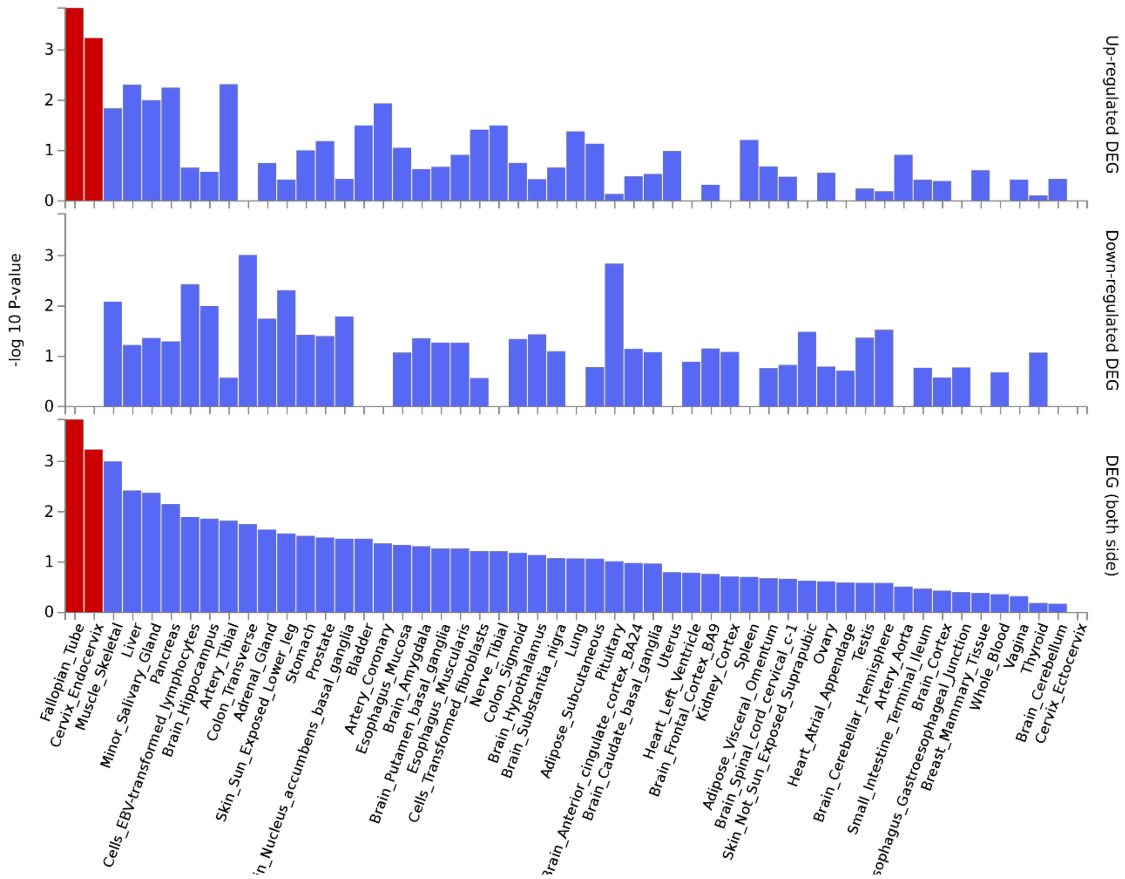

-log 10 P-value
Up-regulated DEG
Down-regulated DEG
DEG (both side)
0
1
2
3
Fallopian_Tube
Cervix_Endocervix
Muscle_Skeletal
Liver
Minor_Salivary_Gland
Pancreas
Cells_EBV-transformed_lymphocytes
Brain_Hippocampus
Artery_Tibial
Colon_Transverse
Adrenal_Gland
Skin_Sun_Exposed_Lower_leg
Stomach
Prostate
in_Nucleus_accumbens_basal_ganglia
Bladder
Artery_Coronary
Esophagus_Mucosa
Brain_Amygdala
Brain_Putamen_basal_ganglia
Esophagus_Muscularis
Cells_Transformed_fibroblasts
Nerve_Tibial
Colon_Sigmoid
Brain_Hypothalamus
Brain_Substantia_nigra
Lung
Adipose_Subcutaneous
Pituitary
Brain_Anterior_cingulate_cortex_BA24
Brain_Caudate_basal_ganglia
Uterus
Heart_Left_Ventricle
Brain_Frontal_Cortex_BA9
Kidney_Cortex
Spleen
Adipose_Visceral_Omentum
Brain_Spinal_cord_cervical_c-1
Skin_Not_Sun_Exposed_Suprapubic
Ovary
Heart_Atrial_Appendage
Testis
Brain_Cerebellar_Hemisphere
Artery_Aorta
Small_Intestine_Terminal_Ileum
Brain_Cortex
sophagus_Gastroesophageal_Junction
Breast_Mammary_Tissue
Whole_Blood
Vagina
Thyroid
Brain_Cerebellum
Cervix_Ectocervix

### *SIGLEC1*

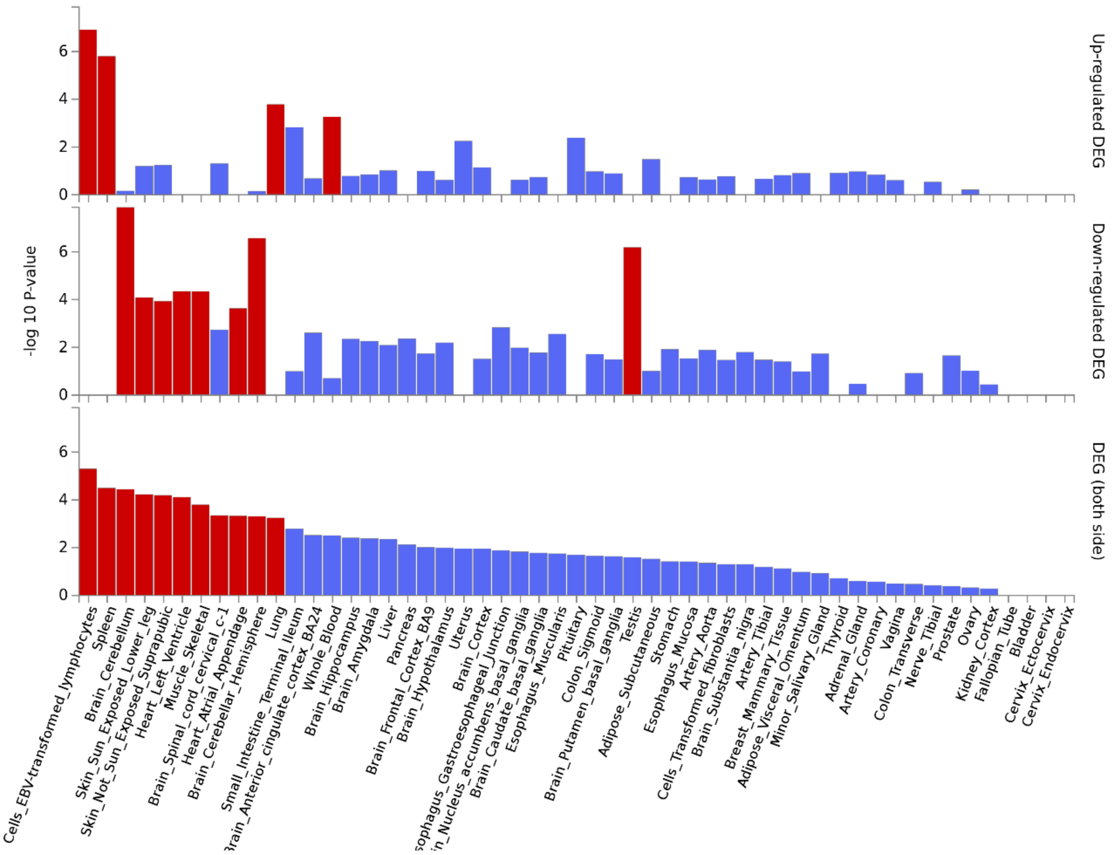

-log 10 P-value
0
2
4
6
Up-regulated DEG
Down-regulated DEG
DEG (both side)
Cells_EBV-transformed_lymphocytes
Spleen
Brain_Cerebellum
Skin_Sun_Exposed_Lower_leg
Skin_Not_Sun_Exposed_Suprapubic
Heart_Left_Ventricle
Muscle_Skeletal
Brain_Spinal_cord_cervical_c-1
Heart_Atrial_Appendage
Brain_Cerebellar_Hemisphere
Lung
Small_Intestine_Terminal_Ileum
rain_Anterior_cingulate_cortex_BA24
Whole_Blood
Brain_Hippocampus
Brain_Amygdala
Liver
Pancreas
Brain_Frontal_Cortex_BA9
Brain_Hypothalamus
Uterus
Brain_Cortex
sophagus_Gastroesophageal_Junction
in_Nucleus_accumbens_basal_ganglia
Brain_Caudate_basal_ganglia
Esophagus_Muscularis
Pituitary
Colon_Sigmoid
Brain_Putamen_basal_ganglia
Testis
Adipose_Subcutaneous
Stomach
Esophagus_Mucosa
Artery_Aorta
Cells_Transformed_fibroblasts
Brain_Substantia_nigra
Artery_Tibial
Breast_Mammary_Tissue
Adipose_Visceral_Omentum
Minor_Salivary_Gland
Thyroid
Adrenal_Gland
Artery_Coronary
Vagina
Colon_Transverse
Nerve_Tibial
Prostate
Ovary
Kidney_Cortex
Fallopian_Tube
Bladder
Cervix_Ectocervix
Cervix_Endocervix
